# Stability of the polarization agent AsymPolPOK in intact and lysed mammalian cells

**DOI:** 10.1101/2024.11.09.622814

**Authors:** Dominique Lagasca, Rupam Ghosh, Yiling Xiao, Kendra K. Frederick

## Abstract

Dynamic nuclear polarization (DNP) solid-state NMR enables detection of proteins at endogenous concentrations in cells through sensitivity enhancement from nitroxide biradical polarization agents. AsymPolPOK, a novel water-soluble asymmetric nitroxide biradical, offers superior sensitivity and faster build-up times compared to existing agents like AMUPol. Here, we characterize AsymPolPOK in mammalian HEK293 cells, examining its cellular distribution, reduction kinetics, and DNP performance. We demonstrate that electroporation achieves uniform cellular delivery of AsymPolPOK, including nuclear permeation, with no cytotoxicity at millimolar concentrations. However, the cellular environment rapidly reduces AsymPolPOK to its monoradical form, with one nitroxide center showing greater reduction resistance than the other. While AsymPolPOK maintains high DNP enhancements and short build-up times in lysates, its performance in intact cells depends critically on delivery method and exposure time to cellular constituents. Electroporation yields higher, more uniform enhancements compared to incubation, but prolonged exposure to the cellular environment diminishes DNP performance in both cases. These findings establish the potential of AsymPolPOK as a polarization agent for in-cell DNP NMR while highlighting the need for developing more bio-resistant polarization agents to further advance cellular structural biology studies.

## INTRODUCTION

Proteins in living cells function within a highly crowded molecular environment, engaging in complex interactions with other proteins, nucleic acids, cofactors, and ligands. These conditions are challenging to replicate in vitro. Doing NMR on proteins inside cells is an appealing approach to studying protein conformation in their native cellular context, preserving the cellular organization, stoichiometry, and interactions of surrounding biomolecules (1-8). Nuclear magnetic resonance (NMR) offers atomic-level resolution of proteins in such complex environments, particularly valuable for examining multiple conformations, cellular interactions, and intrinsically disordered regions (3, 5, 8-11).

Dynamic nuclear polarization (DNP) can dramatically enhanced solid-state NMR sensitivity, enabling detection of proteins at endogenous concentrations (3, 7, 12-16). This technique harnesses the high polarization from stable free radicals, transferring it to target nuclei through microwave irradiation (17-19). The radicals are typically introduced into cellular samples by doping samples with millimolar concentrations of the polarization agents. Cross-effect DNP, which couples two unpaired electrons to one nucleus, has proven particularly efficient using nitroxide biradicals as polarization agents (20). However, the cellular environment poses a significant challenge: the cellular environment reduces nitroxides, diminishing their effectiveness as polarization agents and limiting sensitivity (21-23). Therefore, the stability of polarization agents and their DNP performance inside mammalian cells defines the effective detection limit for proteins inside cells.

Studies examining nitroxide biradical stability under varying cellular conditions began with TOTAPOL, one of the first cross-effect biradical DNP polarization agents (24). In bacterial systems, TOTAPOL demonstrated markedly different reduction kinetics between whole cells and lysates (25). Whole cell pellets rapidly reduced TOTAPOL (2.5 mM) at a rate of 0.18 mM/min, leaving only 6.4% biradical after 10 minutes. In contrast, lysates showed slower reduction (0.043 mM/min), with 64% biradical remaining after the same period. The behavior of AMUPol (26), a widely used water-soluble polarization agent for biomolecular DNP NMR followed a similar pattern. Like TOTAPOL, AMUPol undergoes more rapid reduction in intact cells than in lysates (22, 27), with approximately 50% conversion to the monoradical form occurring before the earliest measurable timepoints (22).

Nitroxyl radicals in five-membered pyrrolidine rings are more biostable than those in six-membered piperidine ring (28). To create a potentially more biostable polarization agent, the piperdine rings in TOTAPOL were replaced five-membered pyrrolidine rings (21). This modification improved the bio-resistance although POPAPOL although the DNP performance was quite modest so despite its improved biostability, its usefulness as a polarization agent for in cell spectroscopy is limited (21).

AsymPolPOK is a highly efficient polarization agent developed through advanced simulations to optimize DNP efficiency and reduce electron spin relaxation (29). This asymmetric nitroxide biradical incorporates a spirocyclohexanolyl-based piperidine radical with phosphate groups that enhance water solubility while preventing aggregation and a pyrrolidine ring for improved rigidity (30, 31). AsymPolPOK demonstrates superior sensitivity at both 9.4 T and 18.8 T, featuring fast buildup times for rapid data acquisition and reduced depolarization compared to AMUPol (29). These properties make it a potentially very appealing molecule for in cell applications because the improved DNP properties – in particular the shorter build up time – will increase the absolute sensitivity and the pyrrolidine ring suggest that this polarization agent may have improved biostability. Indeed, in a recent comparative study, AsymPolPOK had the best DNP performance in buffer with reducing agents and in cell lysates (21). However, its behavior in intact mammalian cells—which possess greater reductive capacity than lysates— remained unexplored.

Finally, the spatial distribution of polarization agents within cells is crucial for interpreting DNP-enhanced NMR data, as sensitivity enhancements depend on proximity between the polarization agent and target nuclei (32, 33). Our previous work with AMUPol demonstrated how delivery methods significantly impact this distribution (27, 34, 35). Electroporation achieved homogeneous distribution throughout cellular compartments, enabling quantitative analysis of the entire structural ensemble. In contrast, incubation resulted in heterogeneous distribution, with lower nuclear concentrations compared to cytoplasmic levels, limiting quantitative interpretation of relative conformational populations. Therefore, characterizing AsymPolPOK’s cellular distribution is essential for proper data interpretation in DNP-enhanced MAS NMR experiments. Its unique molecular features—the different nitroxide centers and phosphate groups—could affect its cellular penetration and compartmental distribution differently than previously studied polarization agents.

To address these gaps, here we characterize the reduction kinetics and DNP performance of AsymPolPOK inside HEK293 cells. We quantify the concentration of AsymPolPOK delivered into cells by electroporation and determine its biostability and cytotoxicity. We then evaluate the distribution and DNP performance of AsymPolPOK when it is delivered to cells by both electroporation and incubation.

## RESULTS

### Reduction of AsymPolPOK in lysates

To determine the rate of reduction of the nitroxide biradical AsymPolPOK in biological environments, we added AsymPolPOK at concentrations relevant for DNP to lysed HEK293 cells and measured the total nitroxide signal intensity by EPR over time (27) (22, 36). The integrated EPR signals report on total concentration, which, in the absence of reduction, is double the AsymPolPOK concentration. Because the two nitroxide radicals of AsymPolPOK can be reduced independently, for accuracy, we report values as the total nitroxide concentration. AsymPolPOK is a stable nitroxide biradical-containing compound. When suspended in buffer, the total nitroxide concentration of AsymPolPOK did not change with time (data not shown). However, when AsymPolPOK was added to mammalian cells, which were then lysed, the total nitroxide concentration of AsymPolPOK decreased in a time-dependent manner for the first several hours and then stabilized. The reduction of EPR signal intensities for AsymPolPOK at concentrations of 10 mM and 20 mM was negligible (**Figure 1A**) but the nitroxide reduction curves for starting AsymPolPOK concentrations of 5 mM and lower were well described by mono-exponential expression with an off-set (*R*^*2*^ = 0.99). After 24 hours of room temperature incubation more than half of the starting nitroxide concentration remained (**Figure 1A**). The reduction rate was proportional to the starting concentration and was 0.043% ± 0.008% of the starting concentration per minute. The offsets of the fits, which represent in the final concentration of nitroxide in the sample when the reduction reaction is complete, were likewise concentration dependent. The dependance of the offset on the initial AsymPolPOK concentration fit to a saturation curve and indicated that HEK293 cell lysates had the capacity to reduce 4.2 mM of nitroxide radical.

**Figure 1.**
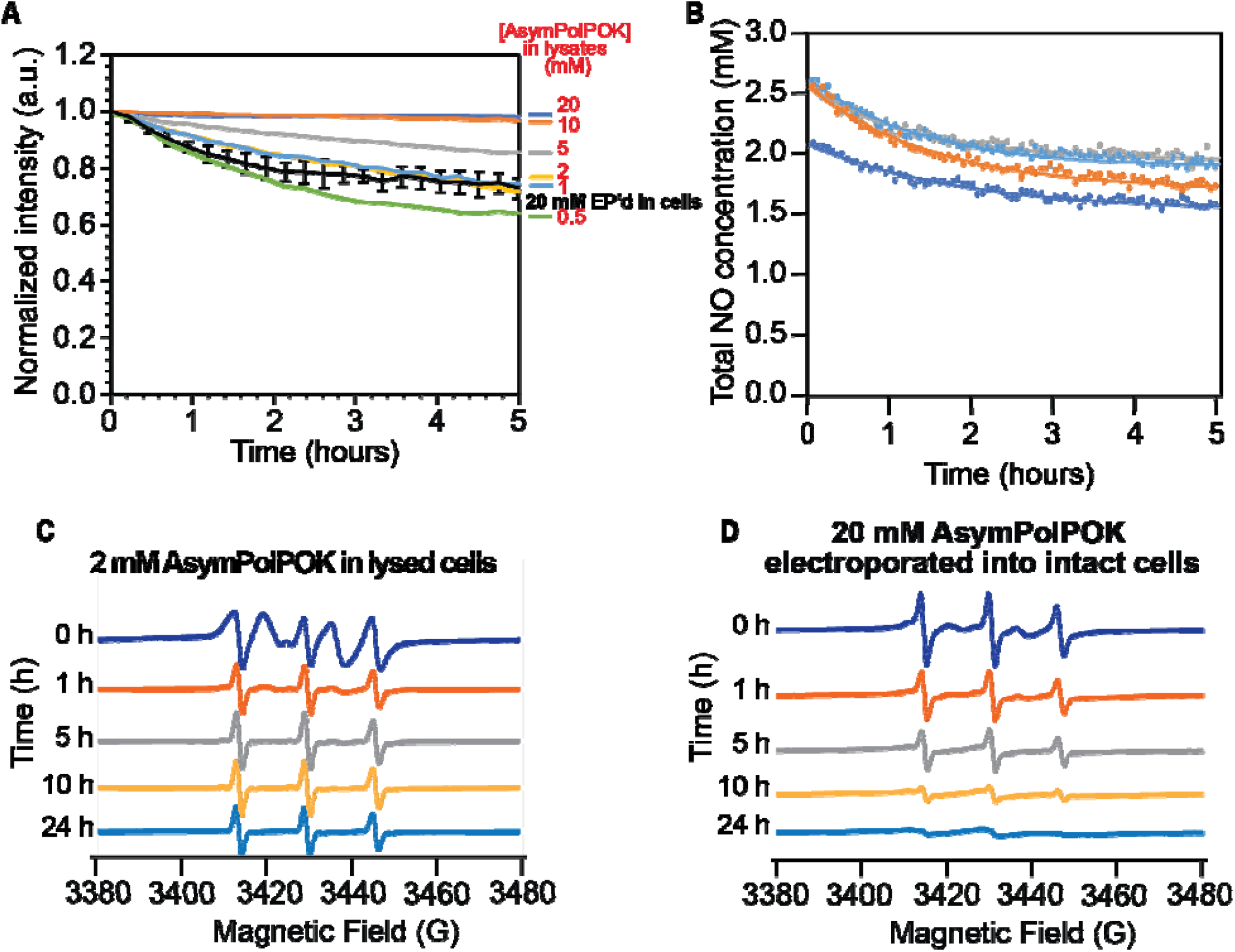
Reduction kinetics of AsymPolPOK in HEK293 cells. (A) Reduction kinetics of AsymPolPOK in cellular lysates. The average AsymPolPOK reduction rate for intact cells (black) is plotted for comparison. Error bars are the standard deviation of 4 replicates from part B. (B) Reduction kinetics of AsymPolPOK inside intact HEK293 cells. AsymPolPOK was introduced by electroporation of cells in the presence of 20 mM AsymPolPOK, followed by a 10-minute room temperature recovery period. After recovery, extracellular AsymPolPOK was removed and EPR measurement began. Total nitroxide (NO) concentration determined by double integration of the EPR spectra is indicated by dots. Lines indicate best fit to an exponential decay. Each biological replicate is plotted in a different color. The concentration of AsymPolPOK delivered to cells was about an order of magnitude less than the concentration present in the buffer at the time of electroporation as expected. (C) EPR spectral changes over the time course of the reduction reaction for cell lysates with 2 mM AsymPolPOK. (D) EPR spectral changes over the time course of the AsymPolPOK reduction reaction in intact electroporated HEK293 cells.

### Reduction of AsymPolPOK inside intact cells

In intact cell samples, we delivered AsymPolPOK to mammalian cells by electroporation and removed extracellular AsymPolPOK before monitoring the decrease in total nitroxide signal intensity over time using EPR (10, 22, 27). Electroporation in the presence of 20 mM AsymPolPOK followed by a 10-minute recovery period and the removal of extracellular AsymPolPOK resulted in a starting concentration of 2.4 ± 0.3 mM (*n* = 4) total nitroxide in the EPR sample volume (**Figure 1B**). As seen with other radicals, electroporation delivered about ∼5% of the AsymPolPOK concentration present in the electroporation buffer to the intact cells. The intact cells were then incubated at room temperature in the EPR spectrometer and the EPR signal was monitored. The rate of reduction of AsymPolPOK was well described by a mono-exponential decay rate (R^2^ = 0.97 ± 0.01), with a reduction rate of 0.041% ± 0.006% of the starting concentration per minute, which is similar to the reduction rate in lysed cells. However, after 24 hours almost no radical remained. This is consistent with a higher reductive capacity for intact cells than for lysed cells.

### The contribution of the DNP-silent monoradical form of AsymPokPOK is more prominent in intact cells than in lysates

While the reduction rates for AsymPolPOK at similar concentrations in lysates and inside intact cells, the shape of the EPR spectra of AsymPolPOK differed. The decrease in radical concentration for both intact and lysed cells was accompanied by changes in the EPR line shape. An isolated nitroxide radical has an EPR spectrum with three peaks. Interactions with other radicals in the sample can broaden or split these peaks. For lysate samples containing 2 mM AsymPolPOK, the initial EPR spectra showed the characteristic splitting pattern of two nitroxide radicals in close proximity (**Figure 1C, 2A** *t* = 0 hrs). Over time, the line shape took on the distinct three-line feature of a nitroxide monoradical (**Figure 1C**, *t* = 5 hrs). In contrast, the EPR spectra even at early time points for intact cells had the strong characteristic features of a nitroxide monoradical (**Figure 1D**, *t* = 0 hrs). With extended incubation, while the EPR line shape ceased to change much, the EPR signal intensity for AsymPolPOK in intact cells continued decrease and was near zero after 24 hours. Thus, intact cells had a greater capacity to reduce AsymPolPOK than lysates.

Next, we determined the individual contributions of the mono- and biradical forms of AsymPolPOK to the EPR spectra. By scaling the spectrum of the biradical form of AsymPolPOK in buffer (**Figure 2A**, blue) to match the intensity of the most down-field and up-field features of the experimental EPR spectra (**Figure 2A**, red), we subtracted the spectrum of the biradical form of AsymPolPOK from the experimental spectrum (**Figure 2A**, black). The corresponding subtracted EPR spectrum contained the characteristic three-peak spectrum (**Figure 2A**). We determined the concentrations of the biradical and monoradical forms of AsymPolPOK individually by double integration. For lysates with 2 mM AsymPolPOK, the biradical form of AsymPolPOK was the major contributor to the EPR spectra at the earliest time points (**Figure 2B**, red) and when fit to a mono-exponential model, the rate of reduction of the biradical form was 0.0048 mM/min. The rate of reduction of the monoradical form to the non-radical form was 0.0035 mM/min, respectively. The accumulation rate of the monoradical form was 0.0065 mM/min. The conversion of monoradicals to nonradicals is slower than the reduction of biradical to monoradical. The biradical form of AsymPolPOK was rapidly reduced to the more stable DNP-inactive monoradical form. Different reduction rate for the biradical and monoradical forms of AsymPolPOK is consistent with two nitroxide radicals having different stabilities in reductive environments.

**Figure 2.**
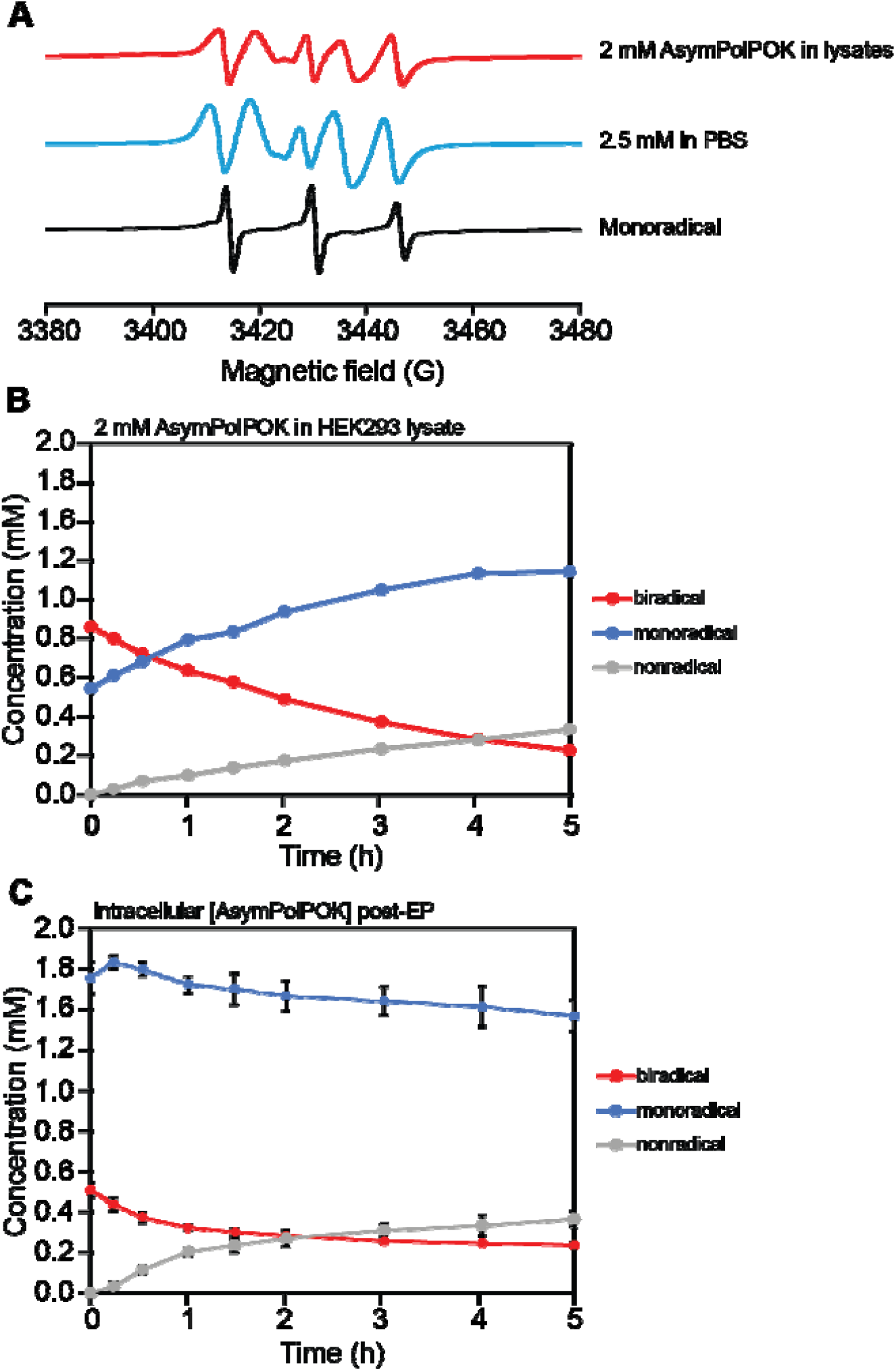
Contributions of the biradical and monoradical populations of AsymPolPOK to the EPR spectra. (A) The concentration (in millimolar) of the biradical (red), monoradical (blue) and completely reduced non-radical forms (gray) changes with time for lysed cells with AsymPolPOK. Error bars indicate the standard deviation of four independent measurements. (B) The concentration of the biradical (red), monoradical (blue) and the completely reduced non-radical forms (gray) for cell lysates after addition of 2 mM of AsymPolPOK. (C) The EPR spectra at t = 0 hr for cells electroporated with AsymPolPOK had a significant contribution from the monoradical form of AsymPolPOK. The subtraction of the EPR spectrum of a completely unreduced sample of AsymPolPOK in the biradical form (red) from the EPR spectrum of cells electroporated with AsymPolPOK (black) results in a spectrum characteristic of an isolated nitroxide radical (blue). Error bars are the standard deviation of 4 replicates.

For AsymPolPOK inside intact cells, where the initial nitroxide radical concentration is ∼2.5 mM, the major form present at earliest time points was the DNP-inactive monoradical form. The DNP-active biradical form of AsymPolPOK comprised about 20% of the population at the earliest time points. The monoradical form remained the major form during extended incubation in the EPR spectrometer (**Figure 2C**). Despite similar starting concentration of radicals to the lysed and intact cells, the starting biradical concentration in intact cells is half of what is found in lysed cells. While the time between initial exposure of the nitroxide biradicals to cellular constituents and measurement is approximately 15 minutes shorter for the lysates than for intact cells (24), this difference in preparation time does not account for the magnitude of difference in the starting biradical concentration. This observation is consistent with a more rapid overall radical reduction rate in intact cells. It also implies that the enhancements of in-cell DNP NMR could be double if AsymPolPOK was protected from inactivation by the cellular constituents.

### Electroporation of AsymPolPOK is non-toxic to HEK293 cells

To determine if exposure to millimolar concentrations of AsymPolPOK is toxic to mammalian cells, we delivered AsymPolPOK to cells by electroporation and allowed the cells to recover for 10 minutes before removing extracellular AsymPolPOK with two wash steps. The cellular integrity was assessed using two complementary approaches: trypan exclusion to assess membrane integrity and regrowth assay to assess toxicity (27). Electroporation of 20 mM AsymPolPOK did not affect the membrane integrity compared to cells electroporated only with buffer (*i*.*e*. a mock electroporation) (10). The trypan exclusion assays for HEK293 cells electroporated only with buffer showed that 96% ± 2% of cells had intact membranes post electroporation (**Figure 3B**). HEK293 cells electroporated in the presence of 20 mM AsymPolPOK had 90 ± 7% of cells with intact membranes post electroporation (**Figure 3B**). Delivery of AsymPolPOK did not alter the membrane integrity of HEK293 cells (unpaired students t-test, *p* > 0.05) and cell viability as assessed by membrane integrity remains high. Next, we assessed the ability of cells to grow after delivery of millimolar concentrations of AsymPolPOK by electroporation to cells. We found that cells exposed to AsymPolPOK had slightly longer lag phase and reached mid-log phase growth a day later than mock-electroporated cells. However, exposure to AsymPokPOK did not affect the cells’ ability to reach confluency (**Figure 3A**). Overall, these findings suggest that AsymPolPOK is not toxic to mammalian cells, making it suitable for applications that require maintaining high cell viability and growth potential.

**Figure 3.**
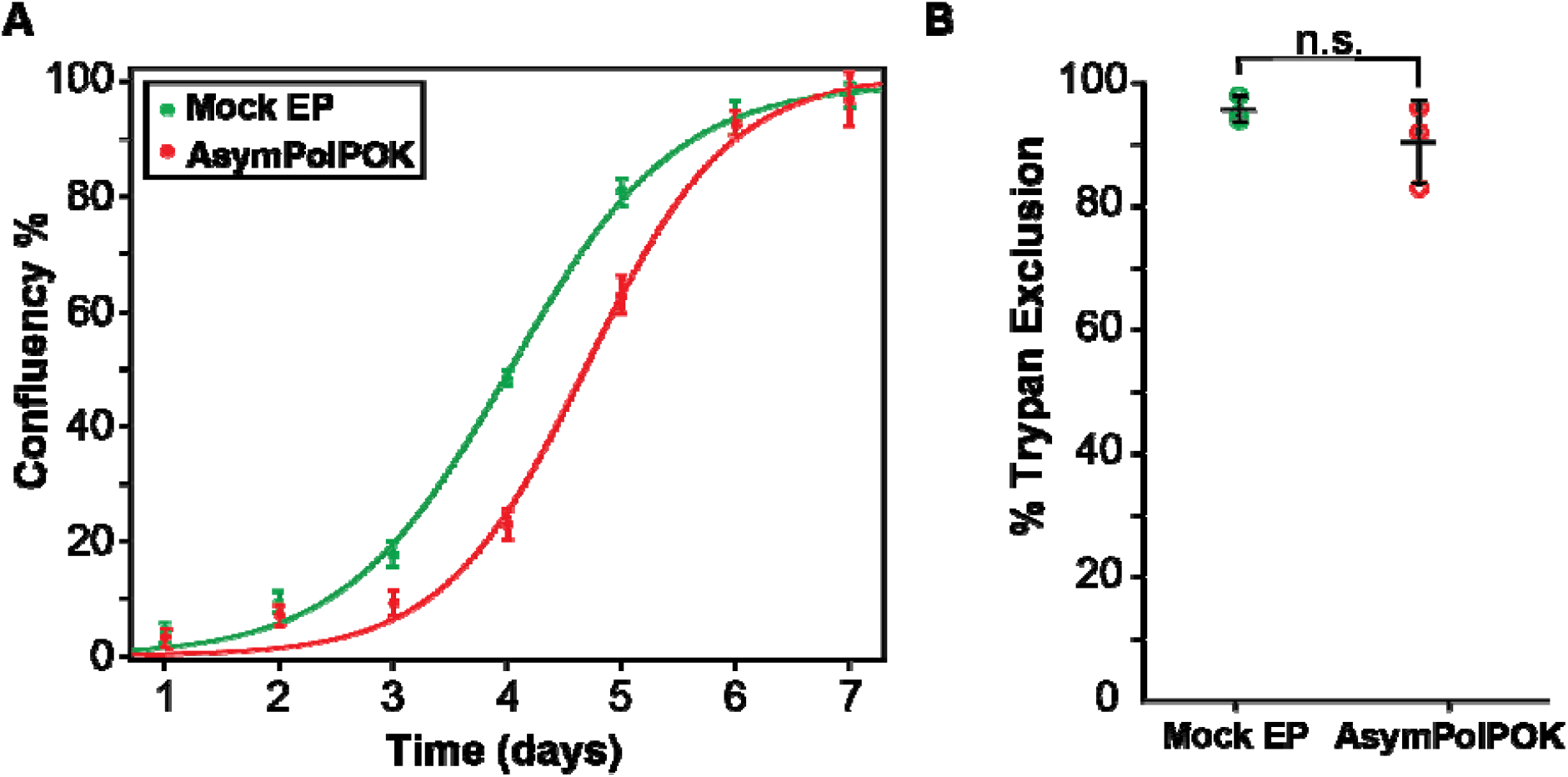
Toxicity assays of AsymPolPOK electroporated in mammalian cells. (A) Growth kinetics as assessed by confluency for cells electroporated only with buffer (green) and AsymPolPOK (red). (B) Percentage of cells with trypan impermeable membranes of cells electroporated only with buffer and AsymPolPOK, colored as in (A) and was analyzed with an unpaired t-test. Bars represent average ± standard deviations; n = 3. No significant difference was observed (n.s., p > 0.05).

### DNP is efficient for all biomass components in cell lysates

We collected ^13^C cross-polarization (CP) spectra with and without microwave irradiation to determine the DNP enhancement for cell lysates with 10 mM AsymPolPOK. We determined DNP enhancements for peaks in the ^13^C CP spectra that are representative of the major biomass components of HEK293 cells; proteins, nucleotides and lipids (27). We found that for cell lysates, the DNP enhancements for proteins, nucleotides and lipids were 93, 87, and 74, respectively. We next assessed DNP build up times (*T*_B,on_) and found that the *T*_B,on_ values for cell lysates for proteins, nucleotides and lipids were 1.7, 1.7, and 2.3 seconds, respectively (**Table 1**). The high enhancements and short build-up times indicate that AsymPolPOK is an efficient polarization agent in cell lysates. To assess the homogeneity of the AsymPolPOK distribution throughout the lysed sample, we examined the β-factor from a stretched exponential fit. For lysed cells with AsymPolPOK the β-factors were 0.87 ± 0.03 across the three biomass components, confirming a homogenous dispersion of AsymPolPOK throughout the sample (**Table 1**) indicating a homogeneous distribution of the polarization agent.

**Table 1.**
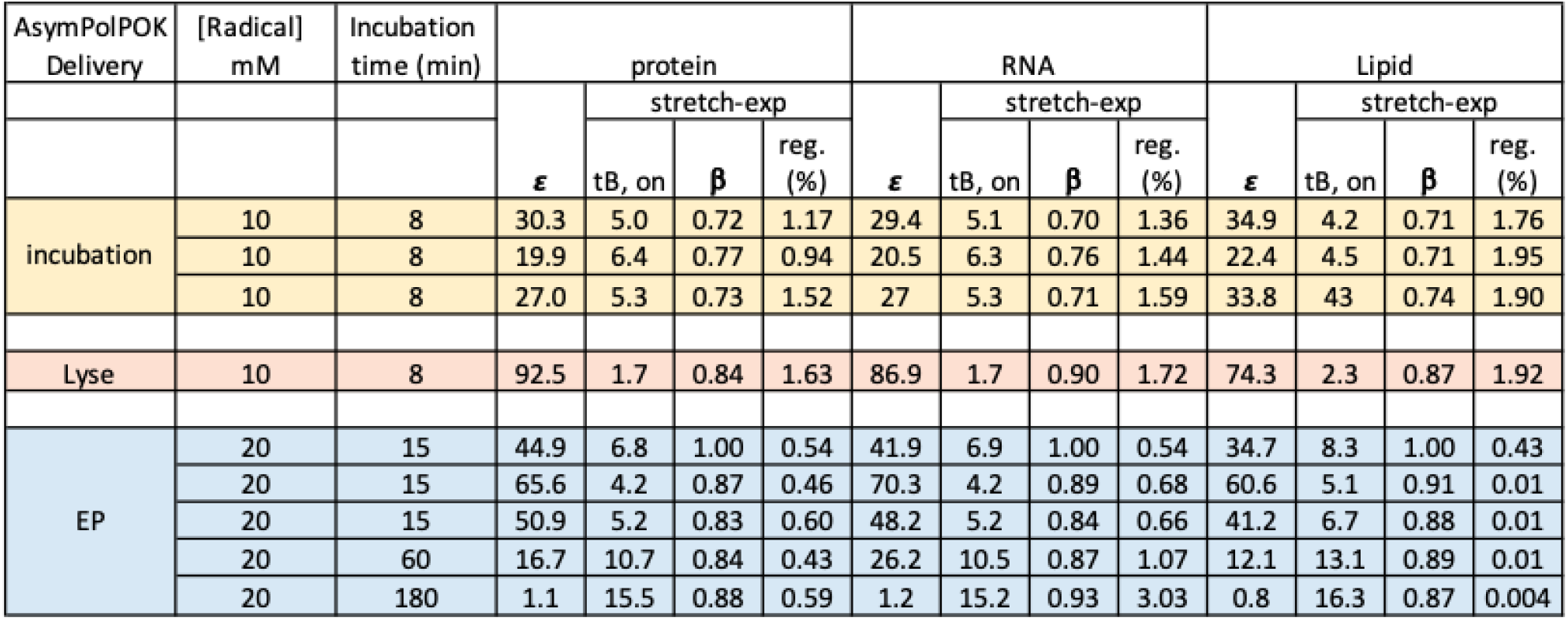
DNP parameters for HEK293 cells that were treated with AsymPolPOK. Stretch-exp report values determined from a fit to a stretched exponential function. T_B,on_ is reported in units of seconds and was determined from 13 time points that ranged from 0.5-60 sec. (0.1, 0.5, 1.0, 1.5, 2.5, 3.0, 3.5, 5.0, 7.5, 10, 15, 30, and 60 sec, respectively). reg. is the regression error, determined as reported in the methods section.

### AsymPolPOK has moderate DNP efficiency in intact cells

We collected ^13^C CP spectra with and without microwave irradiation to determine the DNP enhancement AsymPokPOK inside intact cells. To determine DNP performance of AsymPokPOK inside intact HEK293 cells, we delivered AsymPolPOK to cells in two ways. We delivered AsymPokPOK to cells by electroporation, allowed 10 minutes for recovery, washed the cells twice to remove extracellular AsymPolPOK, and then cryoprotected the cells and collecting for DNP NMR experiments. We also delivered AsymPolPOK by incubating cells in 10 mM AsymPolPOK, cryoprotecting cells and collecting DNP NMR experiments. In cells where AsymPolPOK was delivered by electroporation, DNP enhancements for proteins, nucleotides, and lipids were 54 ± 11, 53 ± 15, and 45 ± 13, respectively (**Figure 4A, Table 1**). In contrast, for cells incubated with AsymPolPOK delivered by incubation, the enhancements were lower: 26 ± 5, 26 ± 5, and 30 ± 7, respectively. We also measured DNP build-up times for these samples. *TB,on* values when AsymPolPOK was delivered by electroporation were 5.4 ± 1.4, 5.5 ± 1.4, and 6.7 ± 1.6 seconds. Interestingly, despite the difference in DNP enhancements the build-up times were similar to those when AsymPolPOK was delivered by incubation, which were 5.6 ± 0.7, 5.6 ± 0.6 and 4.3 ± 0.1 seconds for protein, nucleotide and lipids, respectively (**Table 1**). To assess AsymPolPOK distribution within intact cell samples, we examined the β-factors from a stretched exponential fit. Electroporated samples had β-factors of 0.91 ± 0.07, indicating a homogenous distribution. In contrast, incubated samples had lower β-factors (0.73 ± 0.02), indicative of a heterogenous distribution of AsymPolPOK distribution through the cell.

**Figure 4.**
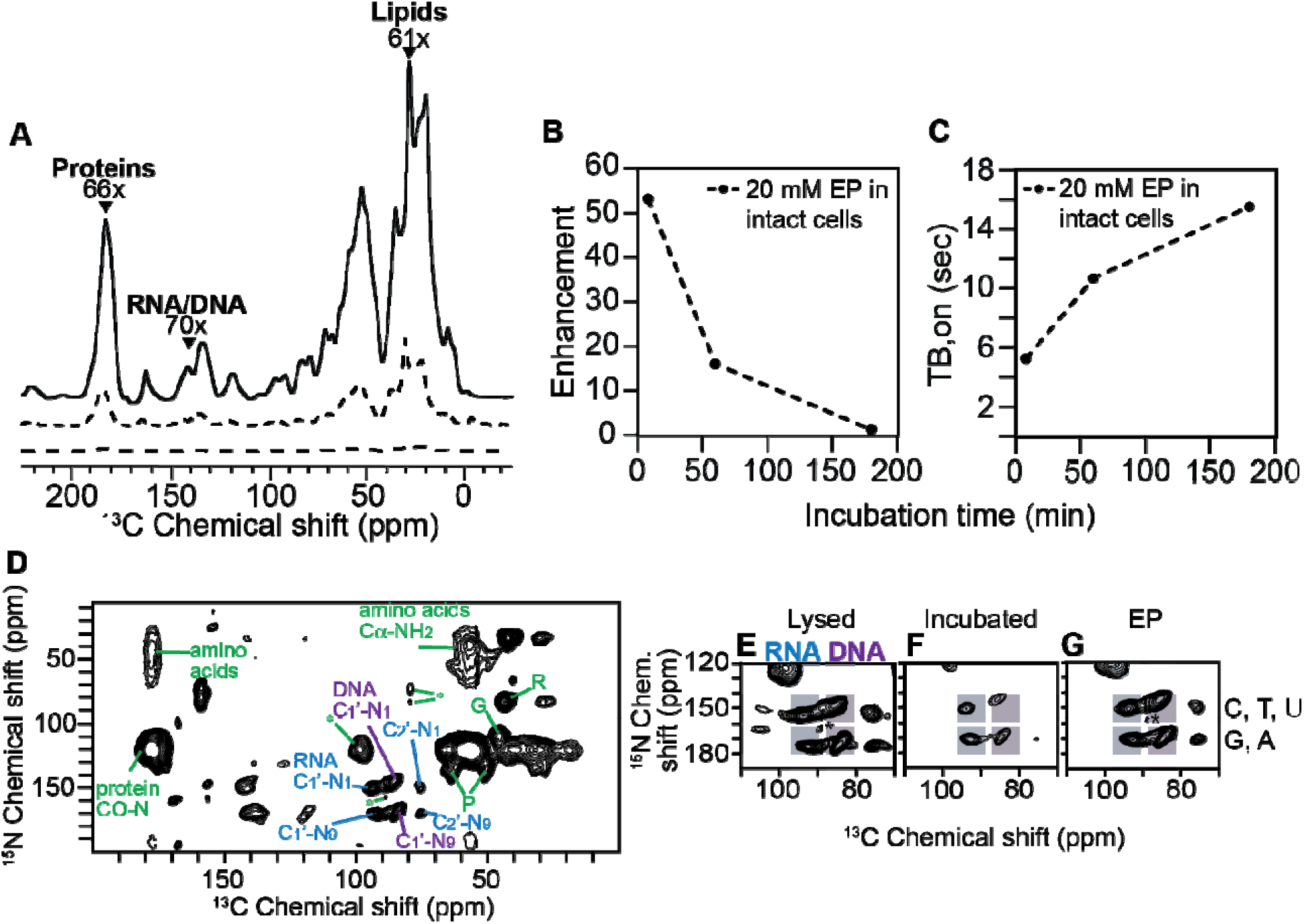
DNP performance of AsymPolPOK in HEK293 cells. A) The DNP performance for cells electroporated with 20 mM AsymPolPOK. Dotted line represents DNP spectra with microwave off and solid line represents DNP spectra with microwave on. Arrows point to major biomass components. B) The DNP enhancement and (C) T_B,on_ performance on intact cell samples electroporated with AsymPolPOK over time. D) 2D heteronuclear correlation spectra (TEDOR) of intact cells electroporated in the presence of 20 mM AsymPolPOK. Selected ^13^C-^15^N correlations from the protein backbone and side-chains (green), from RNA (blue), and from DNA (purple) are annotated. Signals from RNA (blue bar) and DNA (purple bar) are resolved in the 2D TEDOR spectra and are expanded for (E) lysed cells with 10 mM AsymPolPOK, (F) intact cells incubated with 10 mM AsymPolPOK, and (G) intact cells electroporated in the presence of 20 mM AsymPolPOK. Chemical shifts for pyrimidines and purines are indicated by the upper and lower gray bars, respectively. Spinning side bands are annotated by *.

### AMUPol transverses the nuclear envelope of intact cells

While RNA is largely localized to the cytoplasm, DNA is mostly localized to the nucleus. RNA and DNA can be distinguished using ^13^C-^15^N correlation spectroscopy. The ^13^C chemical shift of the C1 carbon of the ribose in RNA molecules is 8 ppm greater than that of the C1 carbon of the deoxyribose in DNA molecules and the ^15^N chemical shift of the N9 nitrogen of the pyrimidine is ∼20 ppm greater than that of the N1 nitrogen of the purine (**Figure 4**). Thus, to determine if AsymPolPOK can enter membrane bound organelles, we collected DNP-enhanced 2D ^13^C-^15^N correlation spectra (TEDOR) on lysed and intact HEK293 cells that were either incubated or electroporated with AsymPolPOK and examined the deoxyribose-purine and pyrimidine regions. We observed strong DNA signals in all the samples (**Figure 4**), indicating that AsymPolPOK was able to enter the nucleus in intact viable cells.

### Reduction by AsymPolPOK cellular biomass degrades DNP performance

Finally, to determine if AsymPolPOK is deactivated by cellular environments at room temperature, we measured DNP enhancements and *T*_B,on_ values for samples prepared with different room temperature delay intervals before freezing. DNP enhancements and *T*_*B,on*_ were measured at 1 and 3 hours post-electroporation. DNP enhancements for protein decreased from 66 to 17 after 1 hour and 1.1 after 3 hours (**Figure 4B, Table 1**), and the build-up times increased from 4.2 seconds to 10.7 and 15.5 seconds (**Figure 4C, Table 1**). These results indicate that extended exposure of AsymPolPOK to the cellular interior decreases DNP performance. Our EPR experiments confirm the deactivation is due reduction of the radical. Thus, minimizing sample preparation time post-AsymPolPOK exposure improves DNP performance.

## DISCUSSION

AsymPolPOK, a highly water-soluble asymmetric nitroxide biradical, exhibits favorable properties for cross-effect DNP at high magnetic fields, including short build-up times that enable faster data collection (29, 37). One of its nitroxide radicals is incorporated in a five-membered ring scaffold, which are known for their biostability. For these reasons, AsymPolPOK may be particularly well suited for DNP-enhanced NMR spectroscopy inside cells but the stability, distribution and toxicity inside cells were unknown. Our characterization of AsymPolPOK in HEK293 cells revealed that while it is not toxic, the biradical is rapidly reduced to a monoradical form in both cell lysates and intact cells. The slow reduction of the monoradical form indicates that the two nitroxide centers have different resistance to reduction by cellular milieu. Notably, the method of delivery significantly impacts AsymPolPOK’s cellular distribution and DNP performance: electroporation achieves homogeneous distribution and high DNP enhancements, whereas incubation results in heterogeneous distribution and more modest enhancements.

In aqueous solutions and cellular lysates, AsymPolPOK demonstrates superior absolute sensitivity compared to AMUPol due to its lower depolarization and shorter build-up time (37). However, in cellular environments, the comparison becomes more complex due to differential radical stability. While one of AsymPolPOK’s radicals – likely the radical in the five membered ring - shows greater reduction resistance than AMUPol’s radicals, its second radical – likely the spirocyclohexanolyl-based piperidine radical with phosphate groups - is more reduction-sensitive. Consequently, at early time points, only 20% of cellular AsymPolPOK remains in the DNP-active biradical form, compared to approximately 50% for AMUPol. The accumulation of AsymPolPOK’s reduction-resistant monoradical form increases overall radical concentration without contributing to sensitivity enhancement.

Despite these differences in radical stability, both polarization agents show remarkably similar DNP performance in cells, with delivery method significantly impacting their effectiveness. With electroporation, both agents achieve homogeneous cellular distribution and comparable DNP parameters: AsymPolPOK (enhancement: 54 ± 11; build-up time: 5.4 ± 1.4 s) and AMUPol (enhancement: 60 ± 6; build-up time: 4.3 ± 0.6 s). Similarly, incubation delivery yields more modest but comparable enhancements for both agents: AsymPolPOK (26 ± 5; 5.6 ± 0.7 s) and AMUPol (30 ± 4; 7.1 ± 0.5 s). While AsymPolPOK outperforms AMUPol in lysates where reduction exposure can be minimized, AMUPol may be preferable for intact cell studies due to AsymPolPOK’s observed effect on cell growth lag phase.

Collectively, this work reveals the complex interplay between molecular design, cellular delivery, and biological stability in determining the effectiveness of DNP polarization agents. While AsymPolPOK’s asymmetric design successfully combines a reduction-resistant radical with enhanced water solubility, its performance in cellular environments highlights the continuing challenge of maintaining DNP efficiency in biological settings. These insights indicate that polarization agent development for in cell DNP must consider the biological stability of both radical centers. Such an approach drove the design of a water solable polarization agent, StaPol-1, which is optimized for high field DNP (38, 39). Such advances will help realize the full potential of DNP-enhanced NMR for studying biomolecular structure and dynamics in cellular environments.

## ACKNOWLEDGEMENTS

D.L. was supported by NIH MB T32 GM008297. R.G was supported by fellowship from the O’Donnell Brain Institute Neural Science Training Program. This work was supported by grants from the National Institute of Health [NS111236, NS134921 and NS124921], the National Science Foundation [1751174], to K.K.F.

## METHODS

### Sample preparation

Samples were prepared as previously described (24). Briefly, HEK293 cells were grown in 10 cm culture dish in complete media containing 10% FBS and 1% Pen-Strep at 37 °C in a humidified incubation containing 5% CO2. At approximately 90% confluency, cells were washed with 1X PBS (pH 7.4) and detached using trypsin. Trypsin was inactivated by the addition of complete media containing 10% FBS and 1% Pen-Strep. The suspended cells were centrifuged at 233 x g for 5 min to pellet the cells. Trypsin and media supernatant were discarded, and the cell pellet was washed with 1X PBS.

Electroporation of HEK293 cells include 50 µL of cell pellet mixed with 50 µL of electroporation buffer containing 40 mM of AsymPolPOK. Cells were mixed with buffer by pipetting the mixture up and down 5 times and electroporated using Lonza electroporator (Lonza 4D-Nucleofactor) using the CM130 pulse program. After electroporation, cells were allowed to recover in the cuvette for 10 min at room temperature and washed twice with 1X PBS. Next, cells were pelleted by centrifugation at 233 x g for 1 min. The supernatant was discarded, and the cell pellet was transferred into a 0.94-mm-inner diameter EPR capillary tube (VWR) and sealed with silicone vacuum sealant.

Cell lysate samples consisted of 50 µL of cell pellet mixed with 50 µL of electroporation buffer containing AsymPolPOK. Cells were lysed via four flash freeze-thaw cycles. Lysed cells were transferred into EPR capillary tube and sealed.

### Cell viability using Trypan Blue dye

10 µL of cells suspended in 1X PBS were mixed with 10 µL of 2X Trypan dye. Total cell viability was measured using automated cell counting machine (Countess II FL, Life Technologies).

### Cellular growth assay

0.2 million cells were plated in 10 cm culture dish containing 10 mL of complete media (DMEM with 10% FBS and 1% Pen-Strep) and allowed to attach overnight. Media was replaced after cells adhered to culture dish and cell growth was monitored for 7-8 days and images were taken. Overall confluency as judged by visual inspection of three fields of view from the tissue culture plate.

### DNP sample preparation

Samples were prepared as previously described (25). Briefly, 50 µL of HEK293 cells were mixed with 50 µL of 40 mM AsymPolPOK in electroporation buffer. Cells were electroporated using the pulse program CM130 in Lonza electroporator, allowed to recover for 10 minutes at room temperature and then washed twice with 1X PBS to remove extracellular biradicals. Cells were then suspended in 50 µL of perdeuterated PBS (85% D2O: 15% H_2_O) and 18 µL of *d8*-glycerol for a final percentage of cryoprotectant in the matrix of 15% (*v/v*). Suspended cells were pelleted into a 3.2 mm sapphire rotor by centrifugation at 233 x g for 2 min and the supernatant was discarded. A silicone plug was inserted into the rotor followed by the ceramic drive tip. The rotor was slow frozen at a rate of 1 °C/min to -80 °C before transferring into liquid nitrogen storage. Frozen samples were cryogenically transferred into the NMR spectrometer (39).

### DNP NMR Spectroscopy

Prepared rotors were transferred in liquid nitrogen directly into the NMR probe that was pre-equilibrated at 100 K. All dynamic nuclear polarization magic angle spinning nuclear magnetic resonance (DNP MAS NMR) experiments were performed on a 600 MHz Bruker Ascend DNP NMR spectrometer/7.2 T Cryogen-free gyrotron magnet (Bruker), equipped with a ^1^H, ^13^C, ^15^N triple-resonance, 3.2 mm low temperature (LT) DNP MAS NMR Bruker probe (600 MHz). The sample temperature was 104 K and the MAS frequency was 12 kHz. The DNP enhancement for the instrumentation set-up for a sample standard of 1.5 mg of uniformly labeled ^13^C, ^15^N proline (Isotech) suspended in 25 µL of 60:30:10 *d8*-glyerol: D_2_O: H_2_O containing 10 mM AsymPolPOK was 67.8 and TBon of 2.4 s.

### EPR spectroscopy

Continuous wave electron paramagnetic resonance (CW EPR) data was collected using a Bruker EMXnano X-band EPR spectrometer operating at a frequency of 9.6 GHz with a center field of 3424 G and sweep width of 172 G. Unless otherwise specified, the following instrumental parameters were used: modulation frequency, 100 kHz; modulation amplitude, 1.00 G; conversion time, 5 ms, microwave power, 0.32 mW; scan time, 5 s. All samples were loaded into 50 µL microcapillary tubes, and their EPR spectra were recorded at room temperature.

The concentrations of the biradical and mono-radical forms of AsymPolPOK were obtained using Bruker’s Xenon software to subtract the spectrum of an AsymPolPOK standard from that of the bi-radical/mono-radical mixture at different time points. For the samples of intact cells (electroporated with AsymPolPOK) and lysate mixed with 1 mM AsymPolPOK, a standard of 0.5 mM AsymPolPOK in water was used for subtraction. For the lysate sample mixed with 10 mM AsymPolPOK, the concentration of the mono-radical contribution to the spectum was determined by double integration and the concentration of standard of 5 mM AsymPolPOK was used for subtraction. For subtraction, the spectra of the standards were shifted, and the gain was adjusted. The concentrations of total nitroxide, bi-radical and the completely reduced forms of the AsymPokPOK were determined by subtraction.

The nitroxide radical reduction curves for electroporated samples and lysates were fit to a mono-exponential model with an offset (y=a*e^(−k*x)+c) using a least squares regression algorithm in MATLAB. In this equation, y is either the total nitroxide concentration in mM or the normalized fractional signal intensity, x is the time, and k is the rate constant. The terms a and c both describe the offset and were fit as independent parameters. The sum of a and c equaled one in our fits.

## Notes

### Competing Interest Statement

The authors have declared no competing interest.

